# Practices for Measuring 3D Organelle Morphology and Generating Surfaces with Amira

**DOI:** 10.1101/2021.09.25.461807

**Authors:** Edgar Garza Lopez, Zer Vue, Prasanna Katti, Kit Neikirk, Michelle Biete, Jacob Lam, Heather K. Beasley, Andrea G. Marshall, Taylor Rodman, Trace Christensen, Jeffrey Salisbury, Larry Vang, Margaret Mungai, Salma AshShareef, Sandra Murray, Jianqiang Shao, Jennifer Streeter, Brian Glancy, Renata O. Pereira, E. Dale Abel, Antentor Hinton

**Affiliations:** Hinton and Garza Lopez Family Consulting Company, Iowa City, IA; Department of Molecular Physiology and Biophysics, Vanderbilt University, Nashville, TN; National Heart, Lung, and Blood Institute, National Institutes of Health, Bethesda, MD; Department of Biology, University of Hawaii at Hilo, Hilo, HI; Department of Internal Medicine, University of Iowa - Carver College of Medicine, Iowa City, IA; Department of Biochemistry, Cancer Biology, Neuroscience and Pharmacology, School of Graduate Studies and Research, Meharry Medical College, Nashville, TN; Microscopy and Cell Analysis Core Facility, Mayo Clinic, Rochester, MN; Department of Biochemistry and Molecular Biology, Mayo Clinic, Rochester, MN; Department of Cell Biology, School of Medicine, University of Pittsburgh, Pittsburgh, PA; Central Microscopy Research Facility, University of Iowa, Iowa City, IA; Fraternal Order of Eagles Diabetes Research Center, Iowa City, IA

**Author notes:** These co-first authors contributed equally (Edgar Garza Lopez, Zer Vue, Prasanna Katti). These authors share senior authorship (Renata O. Pereira, E. Dale Abel, Antentor Hinton). Correspondence: Antentor Hinton, Jr.

**Keywords:** Amira, SBF-SEM, FIB-SEM, Segmentation, Organelles, 3D Reconstruction, 3D Imaging, Mitochondrial Imaging, Volume Analysis

## Abstract

Analysis of 3D structures is of paramount importance in cellular biology. Although light microscopy and transmission electron microscopy (TEM) have remained staples for imaging cellular structures, they lack the ability to image in 3D. However, recent technological advances, such as serial block-face scanning electron microscopy (SBF-SEM) and focused ion beam scanning electron microscopy (FIB-SEM), have allowed researchers to observe cellular ultrastructure in 3D. Here, we propose a standardized protocol using the visualization software Amira to quantify organelle morphologies in 3D; this method allows researchers to produce accurate and reproducible measurements of cellular structure characteristics. We demonstrate this applicability by utilizing SBF-SEM and Amira to quantify mitochondria and endoplasmic reticulum (ER) structures.

**GRAPHICAL ABSTRACT:** 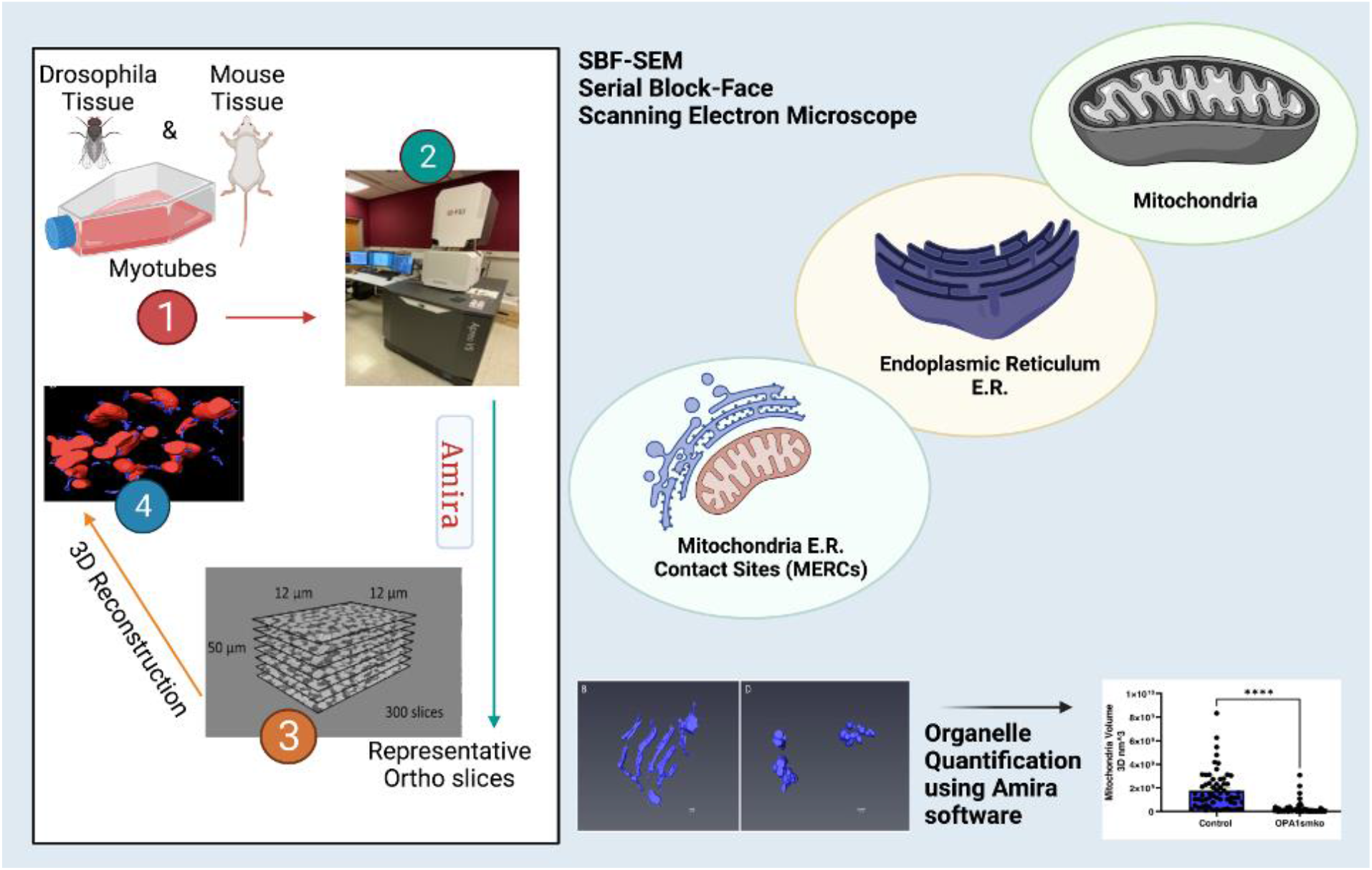

## INTRODUCTION

Cellular organelle research increases understanding of the physiological functions of cells such as vital cellular processes including apoptosis, respiration, and mitosis. Organelles involved in metabolism, such as mitochondria and endoplasmic reticulum (ER), are some of the most studied cellular structures. Because these organelles play major roles in regulating homeostasis and ensuring organism survival, it is important to study their various structure-dependent functions. Mitochondria are typically associated with their role in oxidative phosphorylation, which is crucial for ATP generation, however, their functions extend beyond energetics [1–3]. For example, Dynamin-related protein-1 (DRP-1) regulates mitochondrial fission and associates with early stages of apoptosis [4,5], and changes in mitochondrial ultra-structures are associated with chemical pathways that regulate calcium, potassium, and other biomolecules [6,7]. In addition, the mitochondrial role in maintenance of calcium homeostasis is connected to cellular apoptosis; cellular calcium levels affect ER calcium levels, which in turn regulate mitosis [8]. Further, calcium levels play a role in regulation of the citric acid cycle and calcium signaling associated with apoptosis [9,10]. Given the role of mitochondria in apoptosis, which is a crucial process involved in cancer, mitochondrial research is critical for cancer treatment [3,8–10]. Because mitochondria and ER are crucial for cellular function and survival, their study is relevant to many disciplines and they have potential as targets for pharmaceutical research on neurodegenerative, cancer, and viral disease treatments [2,7].

Imaging of mitochondria and ER has typically been performed using light microscopy and electron microscopy (EM). Despite advances in light microscopy imaging in recent years, EM continues to provide unmatched high resolution ultrastructural detail and 3D visualization [11]. Scanning electron microscopy (SEM) and transmission electron microscopy (TEM) are the most common types of electron microscopy used to study organelle ultrastructure [12–15]. SEM utilizes high-resolution back-scatter detection to provide detailed information of the surface of a sample. TEM, alternatively, produces nanometer-resolution images by transmitting electrons through an ultrathin section of a sample. The novel benefit of volume electron microscopy is the ability to generate near-TEM resolution images using backscatter detection of a block face rather than from an ultrathin section. With an in-chamber ultramicrotome (SBF-SEM) or ion beam (FIB-SEM), samples can thus be continuously sectioned and the block face imaged through very large volumes in the z-plane. The high-resolution stack of images that is generated can in turn provide for unprecedented visualization and analysis of ultrastructure in 3D. FIB-SEM and SBF-SEM are two commonly used methods for generating EM data for 3D reconstruction and both are viable options for the protocol described here. While these are the most common methods, other EM imaging techniques such as automated tape-collecting ultramicrotome scanning electron microscopy (ATUM-SEM) may also be employed [16].

This protocol utilizes SBF-SEM for several reasons. SBF-SEM is a relatively new technique that was developed at the Max Planck Institute in 2004 by Horstmann et al. and has become an established technique to gather volumes of data from various biological samples [17,18]. In SBF-SEM, sectioning is automated and typically involves heavily mordanted samples allowing for backscatter detection and large depths of sectioning of the block face in the z-plane [19,20]. As opposed to some other 3D methods of reconstruction, SBF-SEM can generate thousands of individual images allowing for higher resolution and detection of specific changes in organelle morphology [19,20].

Emerging imaging technologies have facilitated visualization of organelles and tissues resulting in high-resolution 3D reconstruction. In many cases, 2D TEM can be more advantageous than SEM. TEM typically has resolution of less than 50 pm whereas the resolution for SEM images are 0.5 nm; additionally, TEM allows for magnifications nearly 20x higher than SEM [12]. Point counting is traditional used for these 2D micrographs, and is a powerful method for obtaining volume [21,22]. However, such methods are ultimately only estimations while 3D reconstruction offers a more accurate representation of the true subcellular structures. This is especially true for ultrastructure such as nanotunnels. 3D reconstruction is especially important for measurements associated with the folds of the inner mitochondrial membrane of mitochondria known as cristae [7]. Although these folds are typically represented as ribbon-like in a 2D plane, this is an inaccurate representation because cristae are tubular structures that vary in both volume and area [7,23,24]. Recent studies analyzing mitochondria have utilized 3D reconstruction allowing for more accurate representations of calcium stores, spatial distributions of metabolites, and cristae size [25,26]. 3D visualization can be crucial in imaging ER and mitochondria, in addition to other subcellular organelles and multicellular bodies, including blood vessels and organs [25,27–30].

Once high-quality images have been obtained by SBF-SEM or another data acquisition method, they can be reconstructed in 3D using Amira. Amira is a user-friendly application that allows for 2D images, slices known as orthos, to be digitally analyzed, segmented, color-coded, and rendered for 3D reconstruction. There are several programs that can be used for 3D image reconstruction including ImageJ, Microscopy Image Browser (MIB), Arivis, Imaris, Dragonfly, and Ilastik. We choose Amira since it offers abilities that uniquely combine segmentation and 3D reconstruction tools to expedite workflows. Amira allows for a high degree of customizability through an animation creation tool, the ability to assign colors to various organelles, interpolation, compatibility with a wide range of import and export files, and simple workflows involving either manual or semi-automated segmentation [31,32] (Supplemental Figure 1). Furthermore, greater modification is allowed for through scripting interfaces using MATLAB and Python [28]. This is important as deep-learning algorithms on Python are being developed to expedite the workflow of segmentation [33]. Due to this flexibility, Amira provides advantages for both beginners and seasoned researchers over less costly open-source software, such as ImageJ. Prior to using Amira, an adequate computer should be acquired along with an understanding of the potential drawbacks of Amira (see Limitations; Table 6).

**Table 1–5:**
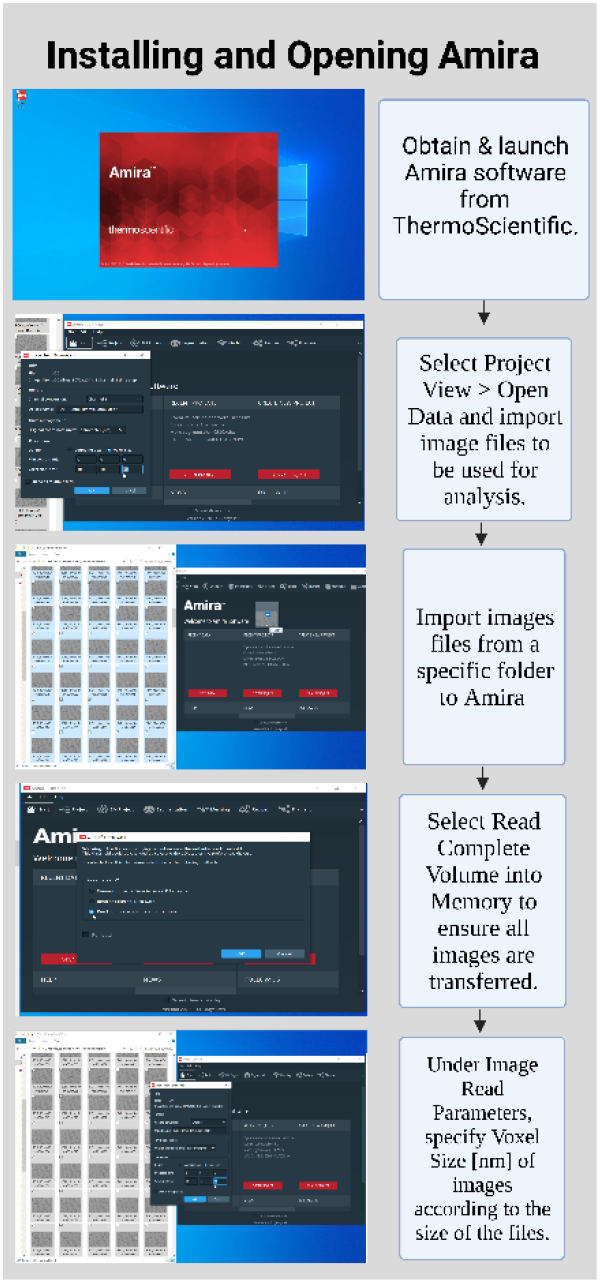
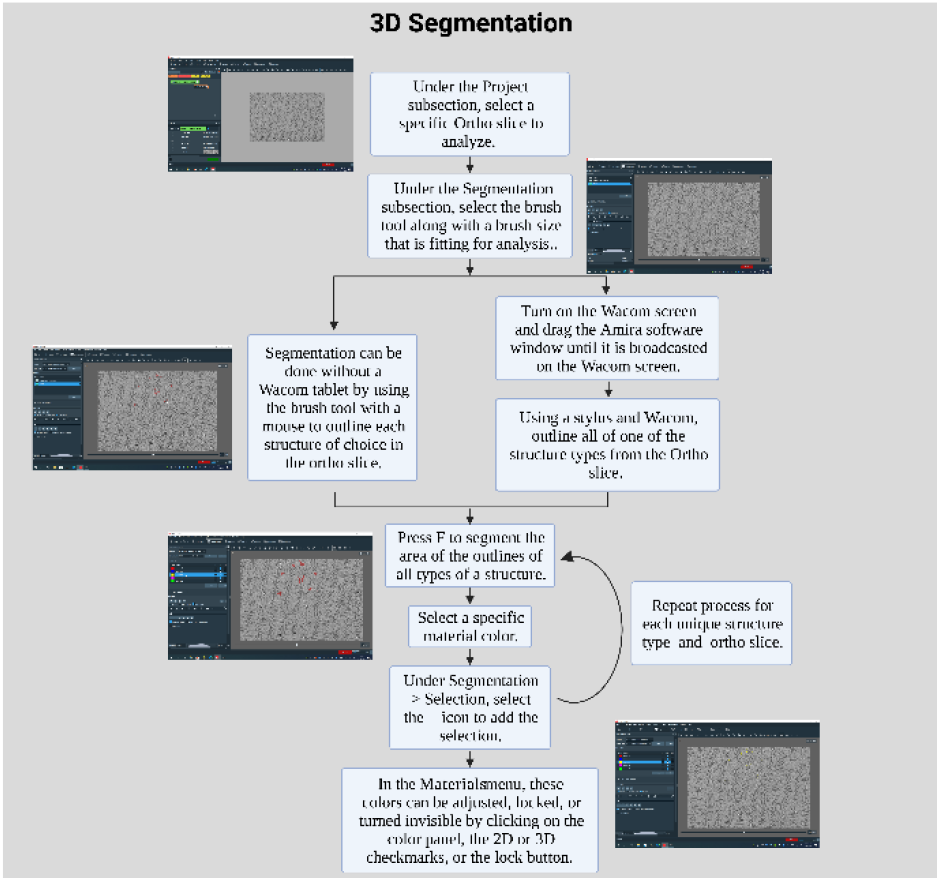
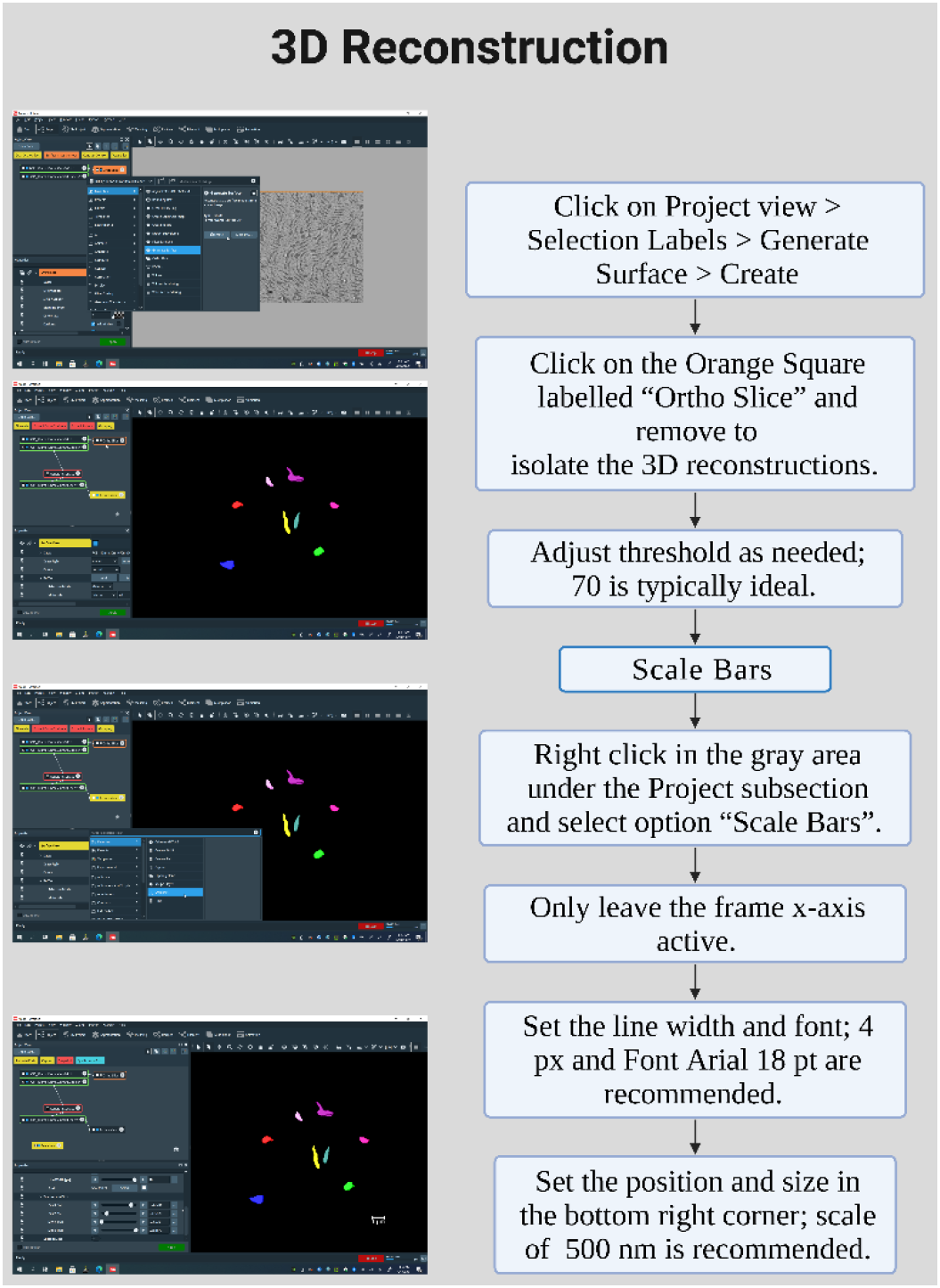
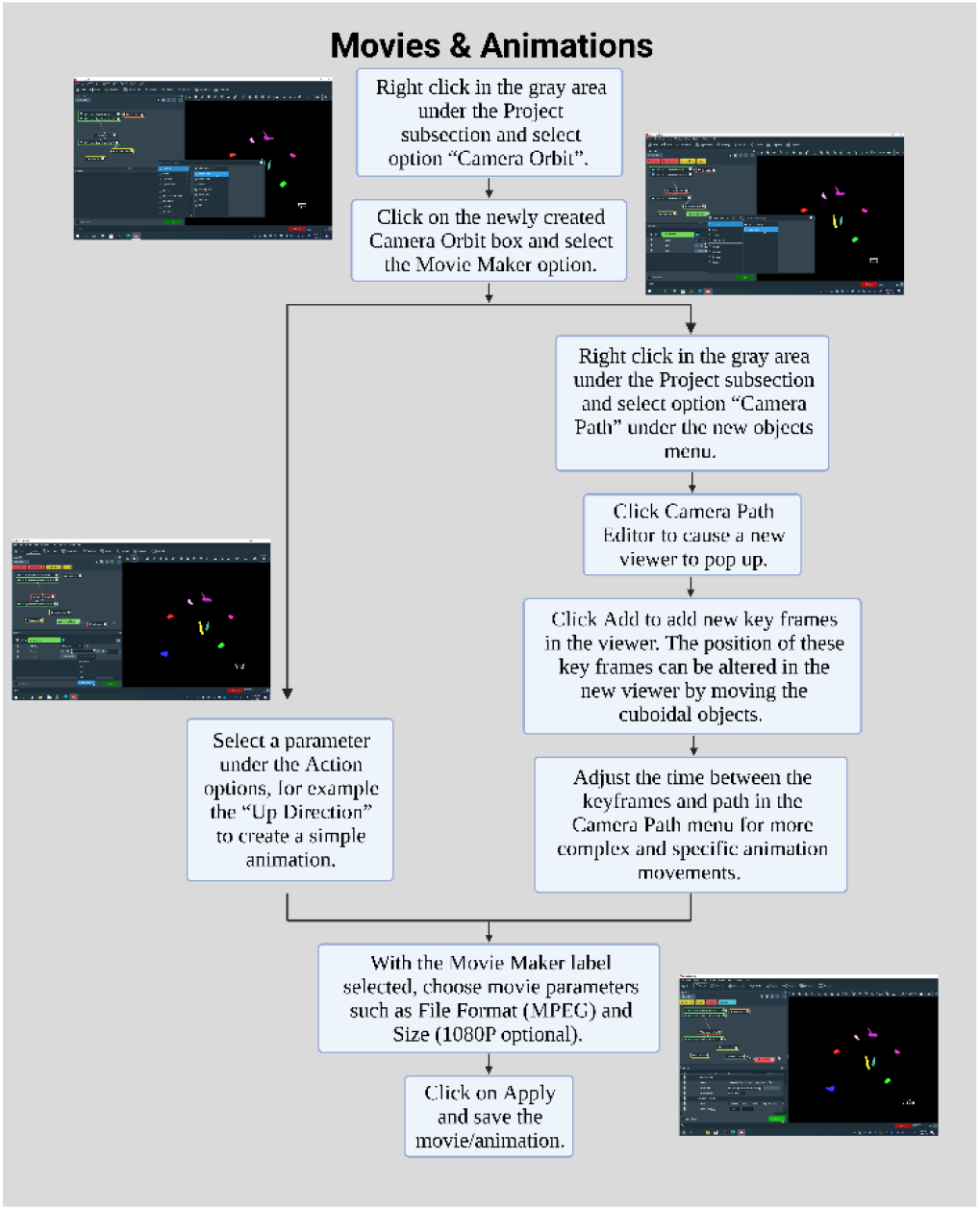
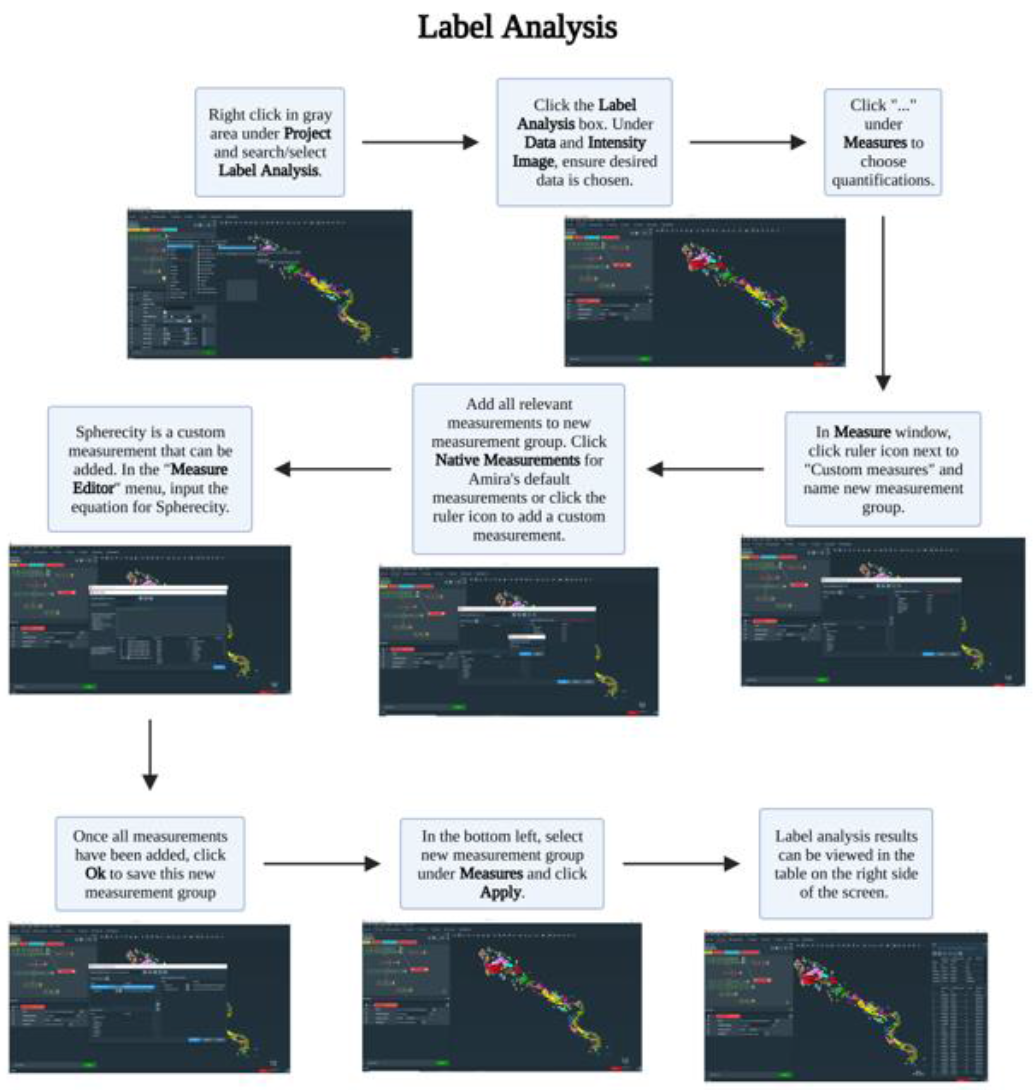
Visualized step-by-step guide for organelle quantification using Amira. Amira offers many customizable options, such as animation styles or semi-automated segmentation, that can be adjusted depending on the objectives and desired workflow. Table 1 demonstrates the basic process to open Amira and import photos into Amira. Table 2 demonstrates the process of using a mouse or Wacom table to manually segment each ortho slice, although this may also be performed with other tools as described in the protocol. Table 3 demonstrates the process of using the segmentation to compile the individual ortho slides into a 3D model of the desired organelles. Table 4 offers a streamlined method for creating videos that can display the 3D models. Methods to create more complex videos are described briefly in the protocol. Table 5 summarizes how to perform quantification using the built in tools on Amira. Although Amira has a high degree of customizability, mastering the software may be difficult. Therefore, this guide provides simple and standardized methods that have many applications and that can be modified or expanded upon as needed.

**Table 6:**
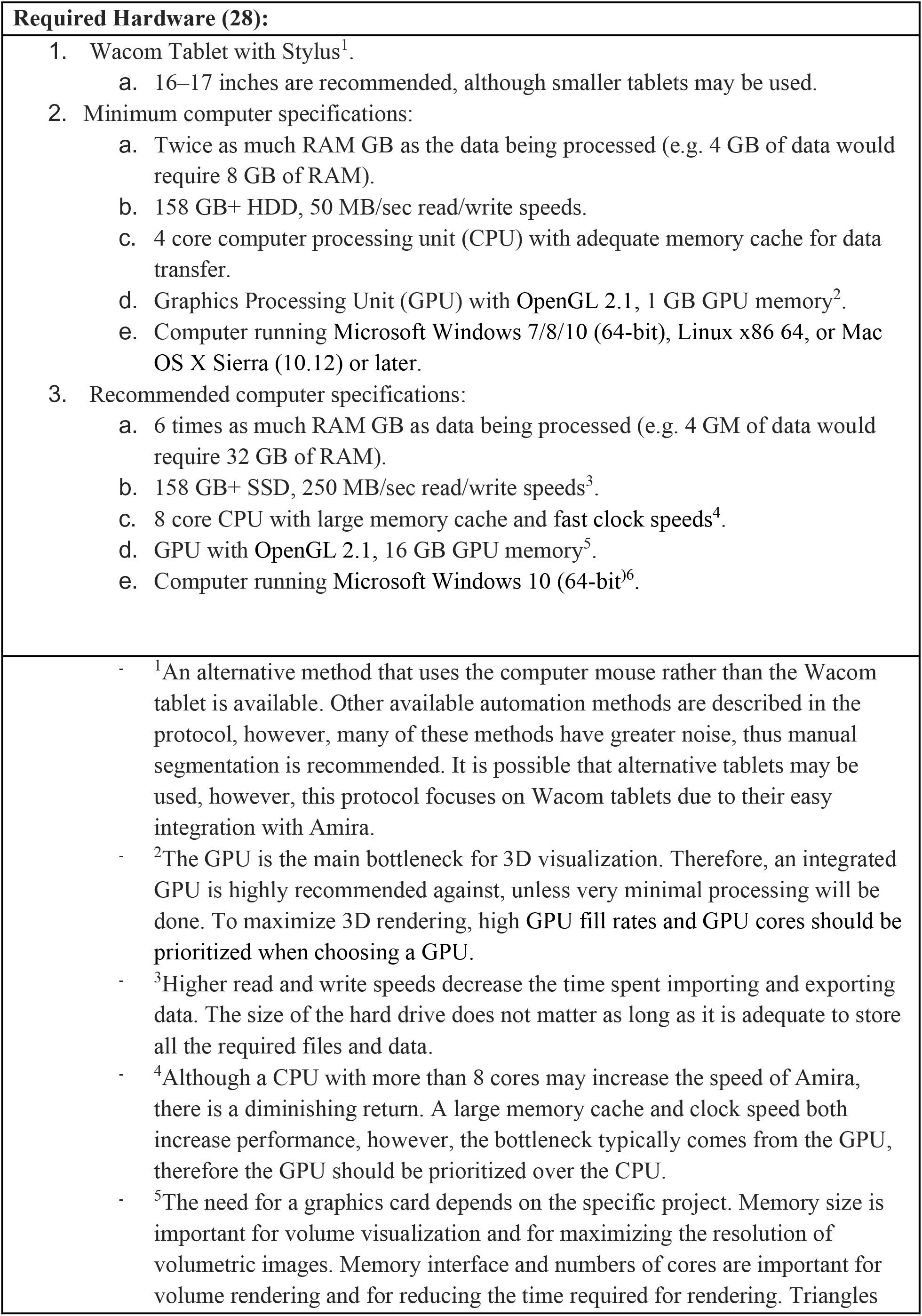
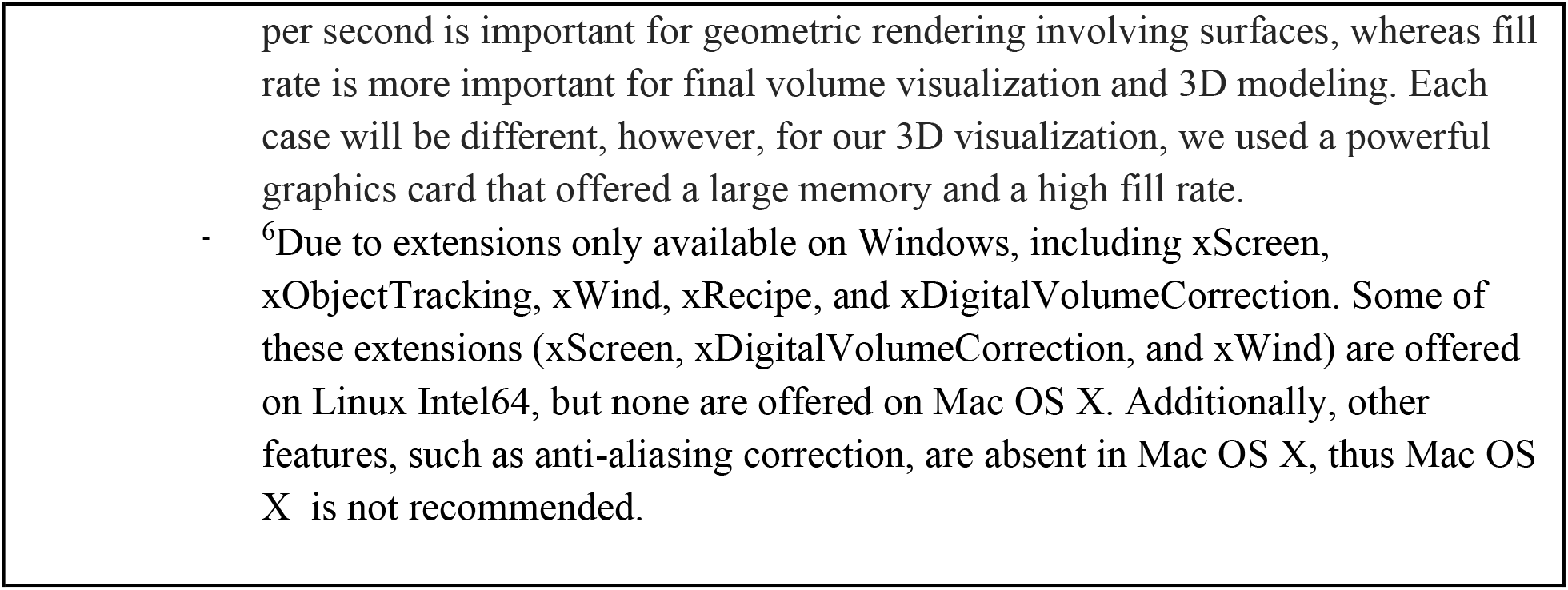
Required hardware for 3D reconstruction of organelles and organs using Amira.

This paper outlines the use of Amira for image segmentation and 3D reconstruction of images acquired through SBF-SEM. In example, these techniques can be used for increasing understanding of structure-dependent functions and the effects of protein mutations on vital cellular components, as well as the potential of these cellular components as therapeutic targets. Here, we outline methods for segmenting and measuring mitochondria and ER volume, as well as the contact sites between these organelles in control conditions versus upon knockdown of optic atrophy-1 (OPA1) and mitofusin-2 (MFN-2).

## PROTOCOL

This protocol is for use on Windows; steps may differ if carried out on another operating system or version of Amira software. These steps have been carried out successfully on other operating systems, however, if issues arise, it is recommended to use the most updated version of Amira on Windows 10.

1. *Installing and Preparing Amira (Table 1)*

1.1. Purchase and download Amira software from ThermoScientific: https://www.thermofisher.com/us/en/home/industrial/electron-microscopy/electron-microscopy-instruments-workflow-solutions/3d-visualization-analysis-software/amira-life-sciences-biomedical.html.
1.2. Open Amira software.
1.3. Select **Project View > Open Data** and import image files to be analyzed from a specific folder.

1.3.1. Select **Read Complete Volume into Memory** to ensure all images are transferred.

***NOTE****: It is important to start with enough material to use the program effectively. Although no minimum number of slices, known as orthos, are required, it is recommended that ∼300 are captured and uploaded. However, not all the images obtained will need to be segmented for the 3D reconstruction. Importing an excess of orthos allows for a select number of them to be used based on quality. It is important that enough orthos, since the structures will not be visible in all of them. Typically, only 50 orthos are ultimately used, but there should be excess to ensure high-quality slices are being chosen*.
1.4. Under **Image Read Parameters**, specify **Voxel Size** (nm) of the images according to the size of the files.
2. *3D Segmentation in Amira (Table 2; Supplementary Figure 1)*

2.1. Under the **Project** subsection, select a specific ortho slice to analyze.
2.2. Under the **Segmentation** subsection, select the **Brush Tool** along with a brush size that is appropriate for analysis; brush sizes 2 and 3 are commonly used. ***NOTE****: This workflow can be sped up by using the **Magic Marker** tool, which allows for automatic segmentation of structures, under Segmentation tools. The specificity can be adjusted in the **Properties** tab on the bottom left and should be adjusted to select only the specific structures to be measured. Pressing shift or control will add or delete, respectively, additional areas when moving the mouse with the Magic Marker tool selected. By checking the **Apply to All Stacks** box, the Magic Marker from a single ortho slice will be extrapolated to all the slices. Although this method is faster, it is less precise and results in more noise*.
2.3. Bring the Wacom device to the computer and turn on the Wacom screen. ***NOTE****: Segmentation can be done without a Wacom tablet by using the brush tool with a mouse; if using this method, proceed from step 2.5 using the mouse cursor rather than the Wacom stylus for tracing. If using a mouse, it is recommended to use the **Auto Trace** tool under Segmentation tools. Although mistakes occur with Auto Trace, it assists in correcting tracing mistakes made when one is performing segmentation with a mouse. Although a mouse is a suitable alternative, we found that a Wacom tablet with a stylus offers much greater precision*.
2.4. Calibrate the Wacom Pen

2.4.1. To calibrate, open the **Wacom Tablet Properties** application on the Wacom tablet.
2.4.2. Select the name of the stylus being used and then select **Calibrate**.
2.4.3. In each corner of the Wacom screen, a cross will appear. Using the stylus, select the middle of the cross.

***CAUTION****: Ensure that the Wacom pen is calibrated. If the pen is improperly calibrated, it will result in the stylus being off, and significant error may be introduced into the segmentation process and the results. For optimal results, it is recommended to recalibrate the pen every 30-60 min; if this is not done, the pen’s accuracy will slowly be lost and will result in a larger offset between the location of the stylus tip and the selection on the screen*.
2.5. Drag the Amira software window until it is broadcasted onto the Wacom screen.
2.6. Using a stylus and Wacom, outline all of one structure type (e.g., mitochondria or blood vessels) in the ortho slice. ***CAUTION****: During this process, it is important to prevent segmentation borders of separate structures from touching. If the borders are touching, Amira will automatically combine the structures together causing individual structures to be falsely merged into one. If the structures of interest are clustered together, several steps can be taken to prevent borders from touching. The brush size may be adjusted down to 1 to lower the width of the brush tool. Alternatively, to avoid merging structures, select the **shrink** option from the **display controls** to reduce the threshold at which Amira combines structures*.

2.6.1. Press **F** to segment the area of the outlined structures once all structures of one type have been segmented.
2.6.2. Select a specific material color for the organelle type. ***CAUTION****: Ensure colors are consistent across all organelles and follow the same pattern. Colors of separate materials cannot be changed to be the same at the end, thus it is important that the colors selected are consistent. It is recommended to segment one organelle type at a time in a consistent color; at the end, the color can be adjusted to the desired color. This avoids the mixing of two separate organelles, which can result in inconsistent 3D reconstructions in the final rendered image. If multiple colors are used, the 3D reconstruction will contain various colors and the analyses, such as 3D volume, will be more difficult or impossible to perform*.
2.6.3. To change the shading of an area, to allow for better visibility, press **D** to change the settings between full color, outline, or gradient. ***NOTE****: It is recommended to use button **D** to change the pattern of an area, especially if a lighter color is being used. This is just a visual change, however, if neglected, structures of interest may be accidentally segmented multiple times, which would waste time and/or cause inaccurate results*.
2.6.4. Under **Segmentation > Selection**, select the **+** icon to add the selection.
2.6.5. Repeat this process for each unique structure type on the ortho slice.
2.6.6. Repeat this process for each ortho slice ensuring that all structure types are consistently the same material color.
2.6.7. Using the **Materials** menu, colors can be adjusted, locked, or made invisible by clicking on the color panel, the 2D or 3D checkmarks, and/or the lock button.
2.7. Segmentation Considerations

2.7.1. This segmentation protocol details the process on an organelle-type basis; however, segmentation may also be performed on an individual-organelle basis (i.e., segmentation of each mitochondrion separately).
  2.7.1.1. To individually segment organelles, repeat Step 2.4 to create new material for each individual organelle as opposed to each type of structure. ***CAUTION****: It is recommended to perform individual segmentation of one organelle type at a time to avoid confusion that may result from working with the different material colors*.
3. *3D Reconstruction of the Segmented Structures in Amira (Table 3)*

3.1. Go to the Project Menu.
3.2. Click on **Selection Labels > Generate Surface**.
3.3. Select **Apply**.
3.4. Rename the newly generated selection cell with the “**. surf**” suffix and click **Surface View**.

3.4.1. The 3D structures should appear over an ortho slice.
3.4.2. Disable the overlaid ortho image. Toggle off to hide the image by clicking on the orange rectangle labelled “**Ortho Slice**” and selecting the **toggle button** (blue square) under the **Properties** menu. ***NOTE****: This is an optimal time to adjust the specific segmented objects that are shown and their colors. While surface view is selected, under **Properties > Materials**, specific Materials can be selected. To remove materials, select **Properties > Buffer > Remove.** This allows for removal of structures that are no longer needed. Alternatively, visibility of materials may be toggled on and off by clicking the blue boxes next to the materials*.
3.5. Make scale bars.

3.5.1. Right click in the gray area under the **Project** subsection and select option “**Scalebars**”.
3.5.2. Leave only the frame x-axis active.
3.5.3. Set the line width and font to something easy to read; 4 px and Arial 18 pt. font is recommended.
3.5.4. Set the position and size to be in the bottom right corner; it is recommended to set the scale to 500 nm.
4. *Creating a Video/Animation in Amira (Table 4)*

4.1. Right click in the gray area under the **Project** subsection and select “**Camera Orbit**”.
4.2. Click on the newly created Camera Orbit box and select **Movie Maker**.
4.3. With Camera Orbit selected, select a parameter under **Action**, such as the “**Up Direction**” to create a simple animation.

4.3.1. Alternatively, one can right click in the gray area under the **Project** subsection and select “**Camera Path**”.
  4.3.1.1. **Camera Path > Camera Path Editor** will cause a new viewer to pop up.
  4.3.1.2. **Camera Path > Add** will add new key frames in the viewer. The positions of the key frames can be altered in the new viewer by moving the cuboidal objects. Add as many keyframes as desired and use the cursor to adjust the path between the keyframes.
  4.3.1.3. Adjust the time between the keyframes in the **Camera Path** menu. This will allow for more complex and specific animation movements.
4.3.2. With the Movie Maker label selected, choose the movie parameters, such as File Format (MPEG) and Size (1080P optional).
4.3.3. Click **Apply** at the bottom left.
4.3.4. Save video.
5. *Performing Quantification (Table 5)*

5.1. 3d length and other length measurements can be set by selecting the ruler icon under the tool icon when viewing the **Project** view.

5.1.1. Ensure a scale has been set in Amira.
5.1.2. Select the **Line** measurement tool and click on the desired surface to measure.
5.1.3. Click on the two points to be measured, for 3d length this should be the Feret’s diameter.
5.1.4. The measurement will be automatically calculated, units may be changed by clicking on the number that appears on the line measurement.
5.2. For other measurements, right click in the gray area under the **Project** subsection and select “**Label Analysis**”.
5.3. Click on the newly created Label Analysis box and ensure properties are visible at the bottom left of the screen.
5.4. Under the **Data** and **Intensity Image** sections, ensure the desired data is chosen for analysis, and ensure for all 3D measurements that “**3D**” is selected under the **Interpretation** section.
5.5. Under the **Measures** section, click on the “**…**” to choose quantifications.
5.6. In the newly opened Measures window, click the ruler icon next to **User Measures**, and name the new group of measurements.
5.7. Add all relevant measurements to the new measurement group.

5.7.1. Area3d, Volume3d, and perimeter measurements, used for quantification in this study, are default measurements on Amira. Select them from the **Native Measurements** box and click the ruler icon in the middle to add to the measurement group.
5.7.2. Sphericity is a custom measurement that can be added in the **Measure Editor** menu that opens when creating a new measurement group. In the output menu, type in the equation for sphericity according to Amira convention: (pi**(1/3) *(6*Volume3d) **(2/3)/Area3d.
5.7.3. Further quantifications are available as default measurements or can be through adding their relevant equation via the method presented in step 5.7.2.
5.8. Once all measurements are added, click the **Ok** button to save this new group for present and future analysis.
5.9. In the bottom left, select the newly created measurement group under the **Measures** section.
5.10. Click the **Apply** button at the bottom left of the properties screen. Data is then able to be copied to graph on the user’s preferred platform, such as GraphPad.

## RESULTS

This protocol provides a method for 3D reconstruction and animation of organelle, sub-cellular, and organ structures. We used this protocol to perform 3D reconstruction of mitochondria and ER.

### SBF-SEM Demonstrates Mitochondrial Changes in OPA1 smKO-Derived Skeletal Muscle

OPA1 is an inner-mitochondrial membrane that regulates mitochondrial dynamics by promoting fusion [4,5,34–36]. Mitochondrial fission involves the splitting of mitochondria and a reduction in volume; whereas, the fusion of mitochondria results in an increase in volume. In addition to maintaining mitochondrial fusion, OPA1 has been found to maintain tight cristae junctions, and regulate apoptosis [4,34,35]. Using 3D reconstruction, the structural dynamics of mitochondrial fusion and fission events, which are usually lost in the 2D plane, may be observed [4,5]. Therefore, we expected to observe structural changes in mitochondria, such as smaller size and reduced connectivity, upon knocking out OPA1. We used high-quality 3D reconstructions of murine skeletal muscle using SBF-SEM (Figure 1A-H, SV1-2) to quantify and measure the mitochondria. Dimensions of the tissue acquired (Figure 1A, E) along with an example ortho slice as a representative image (Figure 1B, F) are both shown. Additionally, 3D reconstruction data is presented in two ways. Ortho slices overlaid with 3D reconstruction shows the representative images of those being reconstructed (Figure 1C, G), while the isolated 3D reconstruction images allow for the making out of finer details (Figure 1D, H). Furthermore, additional details may be understood from the videos, which showcase the mitochondria of each from a variety of angles (SV1-2). Both 3D mitochondrial length and volume were substantially decreased with OPA1 ablation (Figure 1I and 1J) in OPA1 smKO skeletal muscle, compared to the wildtype control. These data suggest that OPA1 produces more fragmented and smaller mitochondria, as observed in 3D, due to increased fission as a consequence of reduced OPA1 levels.

**Figure 1.**
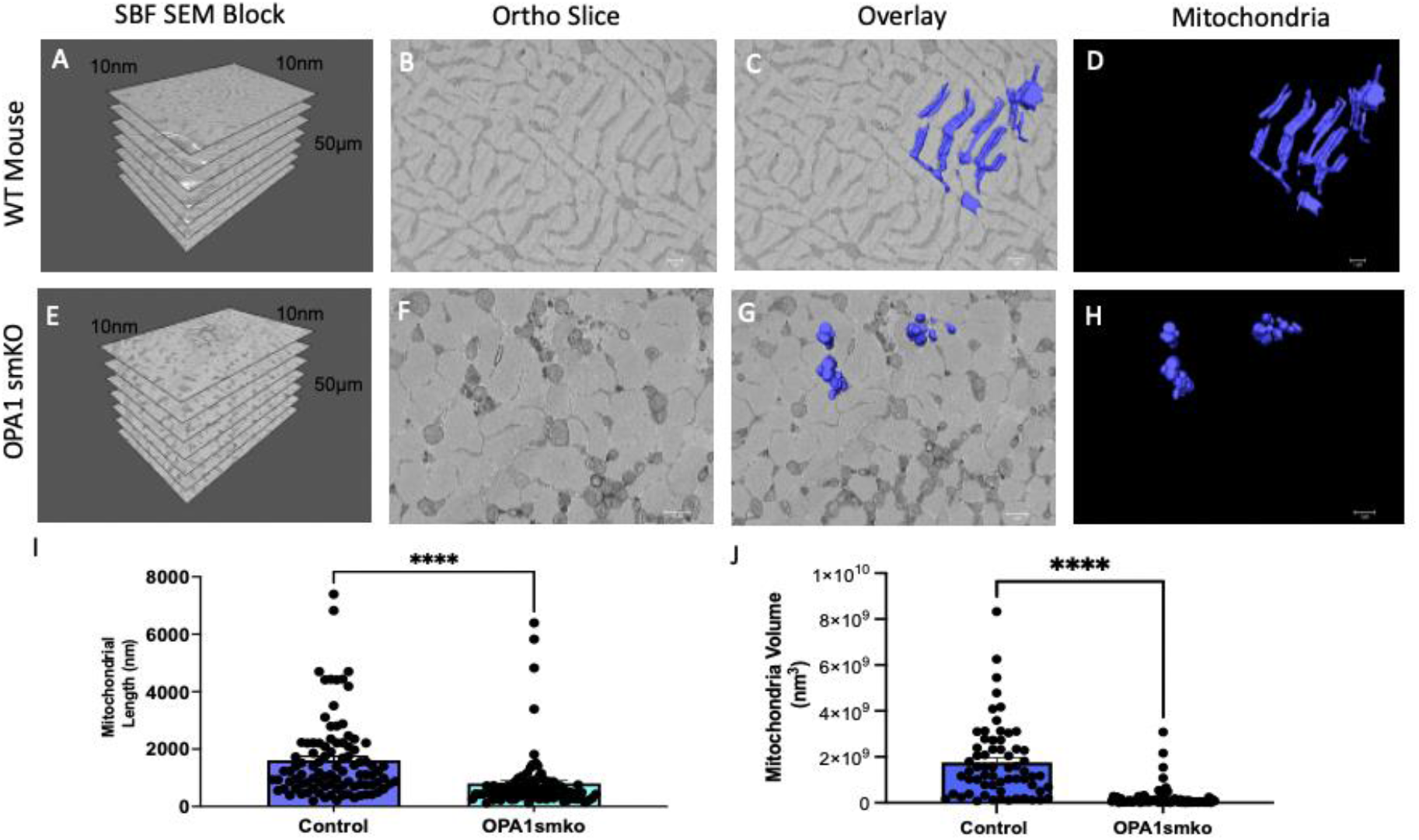
OPA1 deficiency in skeletal muscle leads to changes in mitochondrial morphology in the mouse. The 3D distribution of single continuous and stationary mitochondria (blue), reconstructed from serial block facing-scanning electron microscopy (SBF-SEM) image stacks of *OPA1* skeletal muscle specific knockout (*OPA1* smKO) mouse muscle (A-H). (A) The dimensions of the captured tissue in wild type mouse and (E) OPA1 smKO, (B, F) along with an example ortho slice for each. (C) The overlay of the 3D surface rendering of mitochondria in a wild type mouse, on top of a representative ortho slice and (D) the 3D surface rendering of mitochondria alone. (G) The overlay of the 3D rendering of mitochondria in OPA1 smKO, on top of a representative ortho slice and (H) the 3D surface rendering of mitochondria alone. (I-J) The 3D mitochondrial length and volume decreased (p<0.001) upon OPA1 smKO.

### 3D Reconstruction Allows for Identification of Mitochondria-ER Contact Sites (MERCs)

Both mitochondria and ER have been demonstrated to play crucial roles in cellular processes, such as apoptosis, metabolism, and steady progression through the cell cycle [1–3,37]. However, the specific locations of the MERCs have recently come under scrutiny. MERCs are important because they mediate the transfer of calcium from ER to mitochondria and serve as sites of bio-signaling, among other important functions [38–41]. Calcium signaling and lipid signaling mediated by MERCs are also important for mitochondrial fission and fusion. Additionally, the 3D reconstruction of MERCs may also allow for better understanding of the disease-driven changes in ER-mitochondria communication.

Here, we present qualitative figures that demonstrate the 3D reconstruction of MERCs in *Drosophila*. Figure 2 shows the standardized method for presenting 3D reconstruction data. Figure 2A shows the dimensions of the ortho stacks analyzed. Figure 2B and C shows a representative ortho slice and the overlaid MERCs, respectively. These panels are useful in their descriptions of data acquisition and sampling of base images used for segmentation. The 3D reconstruction showing the mitochondria and ER is useful in highlighting volumetric spaces across which the mitochondria (red) and ER (blue) are in contact (Figure 2D).

**Figure 2:**
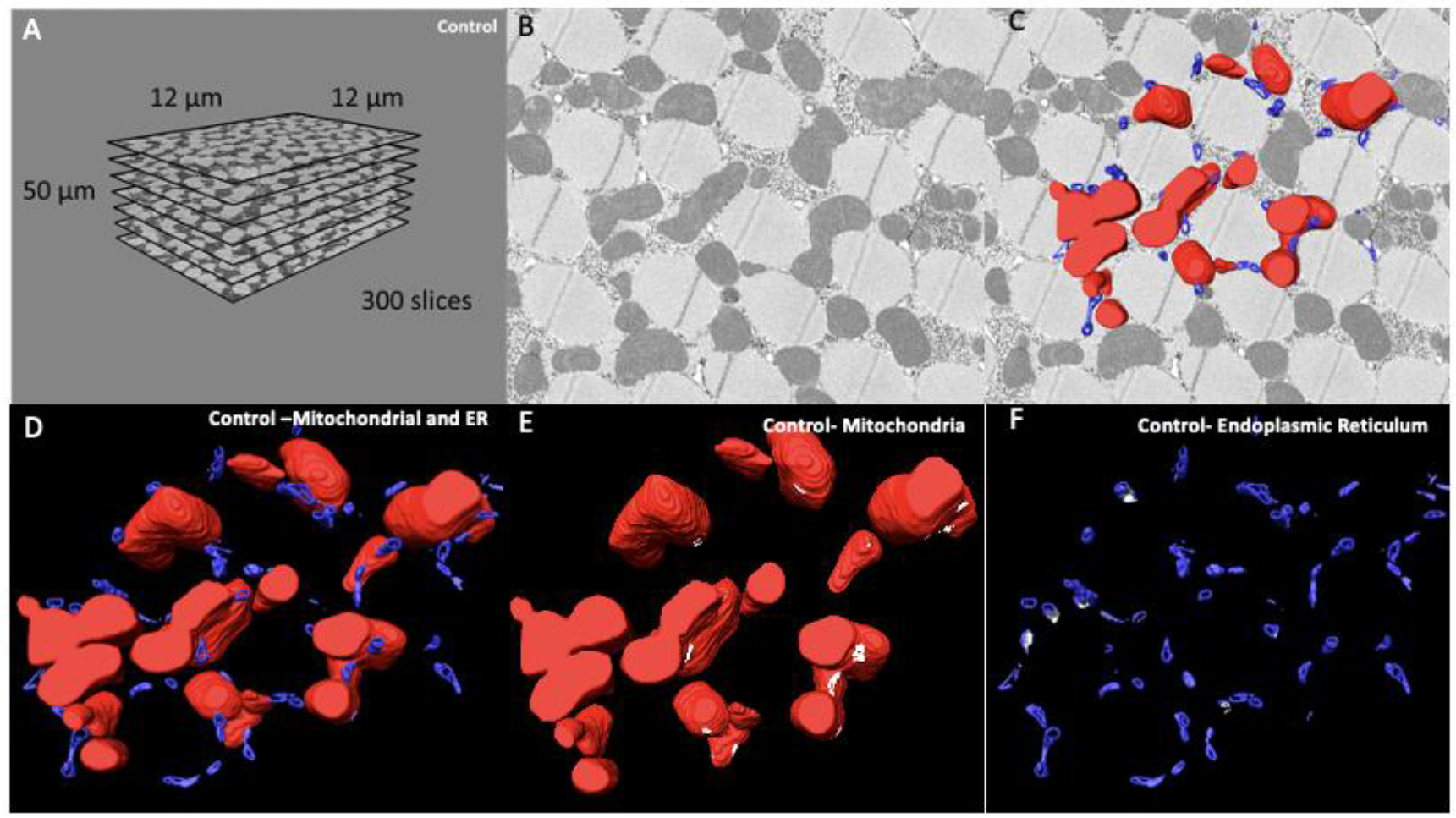
6-panel presentation of 3D reconstruction images and ortho slices. This figure is an example of how to present the ortho slices and the 3D reconstruction images. This example shows 3D reconstruction of several organelles in mouse tissue. (A) On the left, several representative ortho slices are presented. The dimensions and amounts of ortho slices for data acquisition and conversion to 3D models are shown. (B) The raw image of an ortho slice. (C–F) Mitochondria are colored red, ER are colored blue, and MERCs are colored white. These data are best presented in several ways. (C) 3D reconstruction overlaid over the ortho image allows for better visualization of the specific structures in the ortho image that are reconstructed. (D) In contrast, the 3D reconstruction not overlaid on the ortho image allows for better visualization of interactions between the 3D structures. (E, F) Finally, Amira also allows for the graying out of specific structures such that only mitochondria or ER are shown in the 3D reconstruction. This is useful to view otherwise difficult to see areas including MERCs.

An alternative presentation method is via an animation generated in Amira (Video 3). Both presentation forms show the many sites where MERCs exist along with a complete visualization of the two critical organelles, mitochondria, and ER. This flexibility in the presentation of qualitative data is beneficial for studies on genes involved in proper MERC formation, structure, and maintenance. To highlight the MERC sites, they were labelled in white in subsequent panels (Figure 2E, F). Data presentation in the animation style allows for a less cluttered 3D view that shows MERC sites clearly. Furthermore, 3D animation counterparts allow for more detailed depictions of the sizes of MERC sites throughout the depths of mitochondria. (Figure 2E, F, Video 3).

### 3D Reconstruction Shows Organelle Morphology Changes Upon Knockdown of MFN-2

To further validate our method for measuring changes in ER, mitochondria, and MERCs, we knocked down mitofusin-2 (MFN-2). MFN-2 is a GTPase that is necessary for mitochondrial fusion [42]. It does this by serving as a physical tether between the ER and mitochondria, helping to mediate their connections and the calcium exchange between them [38,39,43,44]. As such, loss of MFN-2 has been shown to both cause mitochondrial dysfunction and ER stress [45,46]. Although the exact role MFN-2 has on MERCs is still controversial, many studies have shown that the loss of MFN-2 results in reduced MERC distances [43,44,47–49]. Given that MERCs are important for regular homeostasis in cells, MFN-2 is important for the effective functioning of cells. As the dysfunction of MERCs has been linked to cancers and metabolic syndromes, it is critical to find methods to measure them [47]. We sought to further elucidate the ability of 3D reconstruction to measure MERCs while, additionally, exploring the debated role of MFN-2 on MERCs.

We used high-quality 3D reconstructions of murine skeletal muscle using SBF-SEM to quantify and measure mitochondria and mitochondria. Beyond the representative images showing the dimensions of the tissues cut and an ortho slice, along with the 3D reconstruction overlaid, 3D reconstruction is also shown in a variety of ways (Figure 3A-L). While traditional methods of doing a single color per organelle is effective in many cases (Figure 3D-E, J-K), we also show how the individual mitochondria have changed upon MFN-2 KD through the pseudo coloration of each individual organelle (Figure 3F, L). This allows for multiple ways to view and measure mitochondria and MERCs. To quantify our results, we measured lengths and volume of both (Figure 3M-P). All these metrics decreased, showing that mitochondria became smaller and less connected upon MFN-2 knockdown. Furthermore, we showed that mitochondria sphericity decreased (Figure 3Q). Together, this suggests dysfunction in mitochondria upon lack of fusion regulation. Beyond this, the decreases in MERC length and volume (Figure 3M-N) validates previous studies that have found MFN-2 loss results in reductions in MERC distances [43,44,47–49].

**Figure 3:**
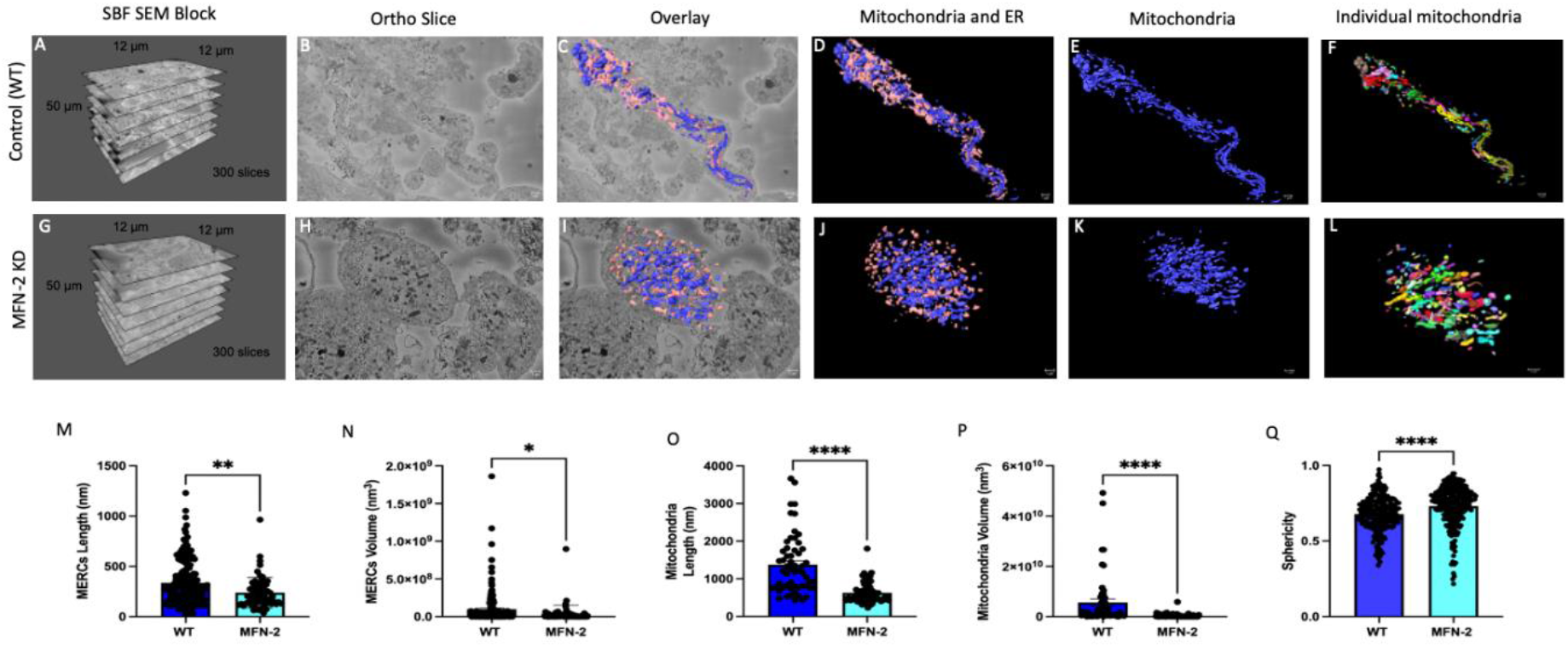
MFN-2 knockdown in mice results in smaller mitochondria and ER, with shorter MERCs. MFN-2 deficiency in skeletal muscle leads to changes in mitochondrial (blue) and ER (pink) morphology in the mice. (A-D) The dimensions of SBF-SEM tissues, isolated ortho slice, 3D reconstruction overlay, and isolated 3D reconstruction in wild type muscle. (E) Wild type muscle 3D reconstructions are also shown with mitochondria colored and (F) ER individually colored. (G-J) The dimensions of SBF-SEM tissues, isolated ortho slice, 3D reconstruction overlay, and isolated 3D reconstruction in *MFN-2* skeletal muscle specific knockdown mouse muscle. (K) 3D reconstructions of mitochondria in a single color and (L) individually colored. (M) When measuring MERCs, MERC length and (P) MERC volume both decreased. Furthermore, the (O) mitochondria length, (P) mitochondrial average volume also decreased. (Q) The sphericity of mitochondria additionally decreased. Significant differences are indicated by asterisks; *, **, and **** indicate *p* ≤ 0.05, *p* ≤ 0.01, and *p* ≤ 0.0001, respectively.

## DISCUSSION

A growing body of cardiovascular and diabetic research demonstrates that ultrastructure imaging is essential to understanding how genes, proteins, and other macromolecules alter organelle morphology [50–52]. However, different methodologies have been used to quantify morphologies of organelle ultrastructure [51–53]. We provide a methodology to use for 3D reconstruction in response to the rise in SBF-SEM use in various biomedical fields. We offer a protocol for Amira software that may be printed for lab use and reference (Supplemental Material 1). This protocol allows for both qualitative and quantitative metrics that may be useful for analysis. However, further exploration and standardization of other 3D reconstruction methods must also be undertaken.

Our results show that 3D reconstruction is an advantageous method to present data involving 3D models. Given that the principal reason for utilizing 3D reconstruction over traditional 2D microscopy methods is the greater level of detail in the 3D plane, the use of animations is important. Animations allow for more details to be visualized than traditional figures (Video 1–5). Our 3D reconstruction animations showcase the complexity of the animations that Amira can produce when key frames, and camera path options are utilized effectively (Video 1–2). For example, when performing the 3D reconstruction of MERCs, the camera panned across all organelles from set angles to allow for identification of all potential MERCs that would go unseen in traditional images (Video 3; Figure 2D–F). Organelles of interest, namely the mitochondria, had key changes elucidated upon OPA1 smKO and MFN-2 KD, with videos providing especially clear views of changes (Figure 1, Figure 3, Video 1-2, 4-5).

The findings presented here demonstrate the power of combining SBF-SEM and Amira software, which can be expanded upon in future research. Qualitatively, MERCs were observed clearly in the animations and the 3D reconstructions of mitochondria and ER (Figure 2E-F, Video 3). The 3D reconstruction of MERCs is useful because it specifies the locations where lipid and calcium homeostasis primarily occur. The 3D reconstructions of mitochondria and MERCs additionally allow for statistical analysis of metrics which cannot be measured as reliably by 2D imaging [21,22] (Figure 3M-Q). As utilized here, quantifications of organelles may be performed upon many conditions, such as gene knockdown. This allows for many future research avenues and results which can only be analyzed properly through EM, and 3D analyses using SBF-SEM and Amira software would be ideal for these studies [23,24,54]. Several other options currently exist for image acquisition that may be used along with or as an alternative to SBF-SEM. In the past, serial section TEM (ssTEM) was the technique of choice due to the relatively simple equipment layout required [51]. However, structural resolution in ssTEM images is limited using heavy metals on the surfaces of structures, and ssTEM is generally unfit for processing large numbers of samples since many methods lack automation, thus requiring human precision, which may reduce reproducibility and reliability [51,52]. SBF-SEM has resolution ranges of less than 10 nm in the x- and y-axis, which is less than the resolution range of less than 1 nm of ssTEM; however, in the z-axis, the resolution is greater for SBF-SEM at less than 50 nm compared to around 60 nm for ssTEM [15,53]. This makes SEM techniques more effective for 3D reconstruction in many cases. FIB-SEM and SBF-SEM are similar in that they both allow for automated imaging; however, FIB-SEM uses an ion beam for cutting and SBF-SEM uses a diamond knife for cutting which allows for greater resolution along the z-axis with FIB-SEM [51]. Although FIB-SEM and SBF-SEM destroy the sample being analyzed, they provide high resolution [51]. Additionally, although typically used in neuroscience, ATUM-SEM is a powerful and automated method that functions by collecting serial sections onto a carbon-coated tape which may later be imaged with SEM [16,23,53]. In general, many ssTEM techniques are quite labor intensive and skill-based, whereas FIB-SEM, SBF-SEM, and ATUM-SEM offer more automatized solutions that allow for high resolution 3D imaging [16,51–53], which dramatically improves interpretation of morphological results of tissue and cell samples. Amira software, and most of the other software options discussed, may be used for FIB-SEM, SBF-SEM, and ATUM-SEM images [55–58]. FIB-SEM and SBF-SEM are quite similar in many ways; the main difference between FIB-SEM and SBF-SEM involves the z-direction resolution of finer details, such as cristae structures and nanotunnels, which are better analyzed by FIB-SEM. However, for studies with large samples, SBF-SEM is faster and offers larger fields of view [53]. Thus, we chose to focus this study on SBF-SEM and we took advantage of the versatility, ease of use, and features of the Amira software for analysis of SBF-SEM images.

Amira offers a variety of useful features designed for the segmentation process, visualization, and quantification of 3D and 4D imaging data, and especially EM data [32]. Amira also offers microscopy-specific image formats for imported files, which makes for seamless processing [32]. Further, this software allows for import of TIFF, JPG, PDF, and DICOM files, whereas some other software packages are more limited in the file types they can import [31]. With a powerful computer, Amira has optimal processing times with previous studies reporting reconstruction of entire organisms in less than thirty minutes [27,59]. Amira is user-friendly and allows researchers to make 3D PDFs of objects using image alignment and color adjustment techniques [59,60]. Amira also provides interactive high-quality volume visualization with orthogonal and oblique slices, volume, and surface rendering, isolines, and isosurfaces for more advanced customization [27]. Following segmentation, Amira allows for post-image processing and analysis, including colocalization, photo bleach correction, and 3D visualization. Thus, Amira can be incredibly simple and user-friendly and can also allow advanced users greater control of statistical analyses by customizing protocols through MATLAB scripts and by outputting data to an Excel format (Figure 1–3 shows statistical analyses methods performed using Amira).

In general, Amira has many quality improvements that reduce the user workload. Amira offers the ability to reduce noise and artifacts in imported images and allows direct manipulation of 3D images for export [61]. Additionally, Amira offers 2D and 3D image filtering to ensure that the full details are shown in the images while, simultaneously, removing the background (See Video 1–5 and Figure 1–3 that showcase the removal of background using Amira). For larger time series data sets, Amira can automate the selection of motion models and custom detection workflows, which can be set to render and detect objects, such as filaments and microtubules, to reduce the overall time required to analyze data [61]. Furthermore, to reduce the stress of large data files, Amira has a novel hybrid file format that reduces the computer space required to perform reconstruction while retaining the original files. Amira also has a failsafe that prevents work from being lost in the event of a software or system crash [61]. As such, Amira is useful throughout the workflow from image segmentation to complete animation of the 3D reconstructed images [31] (See Figure 1 for panel types that can be used in Amira). Amira is highly useful in that it allows for easy-to-obtain images of 3D structures and for more complex custom-shot animations that allow for better examination of structural dynamics [32] (See Video 1–5 and Figure 1–3 which showcase 2D images and 3D reconstruction animations created with Amira). For these reasons, Amira is a powerful tool to use in conjunction with SBF-SEM to generate high-quality 3D reconstructed organelles.

## LIMITATIONS

Although Amira has been used for a range of 3D reconstruction techniques, there are some limitations [27–29]. First, Amira may not be accessible to all researchers due to its cost. Beyond the cost of the software, a high-end computer with a powerful processor and graphics card is necessary to properly process information in a timely fashion (Table 6). Another limitation is the time required to perform manual segmentation. Although tracing structures by hand produces very accurate results, this process can take tens or hundreds of hours depending on the structure of interest, dataset size and number [32]. However, segmentation may be more automated by skillful users. A third limitation is that our method provides only a snapshot of subcellular structures at one point in time. Due to the fixation requirement, our method is unable to document changes in organelle morphology over time. Therefore, other methods, such as organelle staining and live imaging, may be required. Additionally, SBF-SEM cannot be repeated on the same sample because the sample will be destroyed as it is segmented. Although SBF-SEM offers quick data acquisition, the sample cannot be reused for future image acquisition. Despite these limitations, our method provides comprehensive quantification of organelle morphology in 3D.

### Online Methods

Online Methods:
| Reagent or Resource | Source | Identifier |
| --- | --- | --- |
| **Chemicals, peptides, and recombinant proteins** |  |  |
| 2% glutaraldehyde in 0.1 M cacodylate buffer | N/A | N/A |
| 3% potassium ferrocyanide | Sigma | Cat# P3289 |
| 0.1% thiocarbohydrazide | Electron Microscopy Sciences | Cat# 21900 |
| 2% osmium tetroxide | Electron Microscopy<br>Sciences | Cat# 19112 |
| 1% uranyl acetate | Electron Microscopy<br>Sciences | Cat# 22400-2 |
| 0.6% lead aspartate solution | MP Biomedicals | Cat# 155180 |
| HPLC grade acetone | N/A | N/A |
| Epoxy 812 hard resin | Electron Microscopy<br>Sciences | Cat# 14900 |
| **Experimental Models: Organisms/strains** |  |  |
| Mouse: C57B16 | Mayo Clinic | N/A |
| *Drosophila: Mef2 Gal4* | VDRC (Vienna) *Drosophila*<br>stock center and Bloomington<br>*Drosophila* stock center. | BS# 27390 |
| *Drosophila: W<sup>1118</sup>* | VDRC (Vienna) *Drosophila*<br>stock center and Bloomington<br>*Drosophila* stock center. | N/A |
| *Drosophila: Opa-1-like* | VDRC (Vienna) *Drosophila*<br>stock center and Bloomington<br>*Drosophila* stock center. | N/A |
| **Software and Algorithms** |  |  |
| Amira Software | ThermoScientific/Amira [61] | RRID: SCR_007353 |
| Image J | Schneider et al. (2012) [62] | https://imagej.nih.gov/ij/ |
| Ilastik | Berg et al. (2019) [63] | https://www.ilastik.org/publications.html |

## RESOURCE AVAILABILITY

### Lead Contact

Further information and requests for resources and reagents should be directed to and will be fulfilled by the lead contact, Dr. Antentor Hinton Jr.

### Materials Availability

This study did not generate any new, unique reagents.

### Data and code availability

Any additional information required to re-analyze the data reported in this paper is available from the lead contact upon request.

## EXPERIMENTAL MODEL AND SUBJECT DETAILS

### Animal Care and Tissue Isolation

A total of 8 male C57Bl6 mice at 12 weeks of age were used to analyze the impact of OPA1 knockout on mitochondrial structure and networking (n = 3-5 per group). Mice were under a 12:12 light: dark cycle with *ad libitum* access to standard chow and water. All procedures were performed using humane and ethical protocols approved by the University of Iowa Institutional Animal Care and Use Committee in accordance with the National Institute of Health Guide for the Care and Use of Laboratory Animals. Mice were anesthetized using a mixture of 5% isoflurane/oxygen and then the gastrocnemius muscle was excised and cut in 1 mm^3^ pieces before proceeding to the SBF-SEM protocol.

### Fly Strains and Genetics

Genetic crosses were performed on yeast corn medium at 24 °C unless otherwise stated. *Drosophila* strain *W^1118^* was used as the control. *Mef2-Gal4* (III) was used to drive the muscle-specific *Opa-11-like* (OPA1) knockdown (KD). *Tub-Gal80^ts^* and *Mef2 Gal4* (BS# 27390) were used for the conditional muscle-specific *Opa-1-like* KD. Genetic crosses were set up at 18 °C and then shifted to 29 °C at the larval stage. *UAS-mito-GFP* (II chromosome) was used to visualize mitochondria. Stocks were obtained from the VDRC (Vienna) *Drosophila* stock center and the Bloomington *Drosophila* stock center. All chromosomes and gene symbols are as described in Flybase (http://flybase.org).

### Serial Block Facing-Scanning Electron Microscopy (SBF-SEM) Processing of *Drosophila* Muscle Fibers

Tissues were fixed with 2% glutaraldehyde in 0.1 M cacodylate buffer and processed using a heavy metal protocol adapted from a previously published protocol [14,64]. Tissues were immersed in 3% potassium ferrocyanide and 2% osmium tetroxide for 1 h at 4 °C, then treated with filtered 0.1% thiocarbohydrazide for 20 min, 2% osmium tetroxide for 30 min, and left overnight in 1% uranyl acetate at 4 °C; several de-ionized H_2_O washes were performed between each step. The next day, samples were immersed in a 0.6% lead aspartate solution for 30 min at 60 °C and dehydrated in graded acetone (as described for TEM). Tissues were impregnated with epoxy Taab 812 hard resin, embedded in fresh resin, and polymerized at 60 °C for 36–48 h. After polymerization, blocks were sectioned for TEM to identify areas of interest, trimmed to 0.5 mm × 0.5 mm, and glued to aluminum pins. Pins were placed into an FEI/Thermo Scientific Volumescope 2 SEM, a state-of-the-art SBF imaging system. For 3D EM reconstruction, thin (0.09 µm) serial sections, 300–400 per block, were obtained from the blocks that were processed for conventional TEM. Serial sections were collected onto formvar coated slot grids (Pella, Redding CA), stained, and imaged as described above. Segmentation of SBF-SEM reconstructions was performed by manually tracing structural features on sequential slices of micrograph blocks. Images were collected from 30–50 serial sections that were then stacked, aligned, and visualized using Amira to make videos and quantify volumetric structures.

### Data Analysis

Data were presented as the mean of the independent experiments indicated; experiments were performed at least three independent times with similar outcomes. Results were presented as mean ± standard error with individual data points shown. Data were analyzed using an unpaired T-test. If more than two groups were compared, one-way analysis of variance (ANOVA) was performed, and significance was assessed using Fisher’s protected least significance difference test. GraphPad PRISM and Statplus software packages were used for t-tests and ANOVA analyses (SAS Institute: Cary, NC, USA). For all statistical analyses, p < 0.05 indicated a significant difference. Higher degrees of statistical significance (**, ***, ****) were defined as p < 0.01, p < 0.001, and p < 0.0001, respectively.

## ACKNOWLEDGEMENTS

This work was supported by NIH grants R01HL108379 and R01DK092065 to E.D.A; Burroughs Wellcome Fund Award Career Awards at the Scientific Interface (CASI), UNCF/BMS EE Just Faculty Fund, the Ford Foundation, and NIH SRP subaward to #5R25HL106365-12 from the NIH PRIDE Program to A.H.J; and 2U54CA 1633069-07 from BCI and R25 GM059994/GM/NIHGMS NIH to H.K.B.

---

**Supplemental Material 1:**
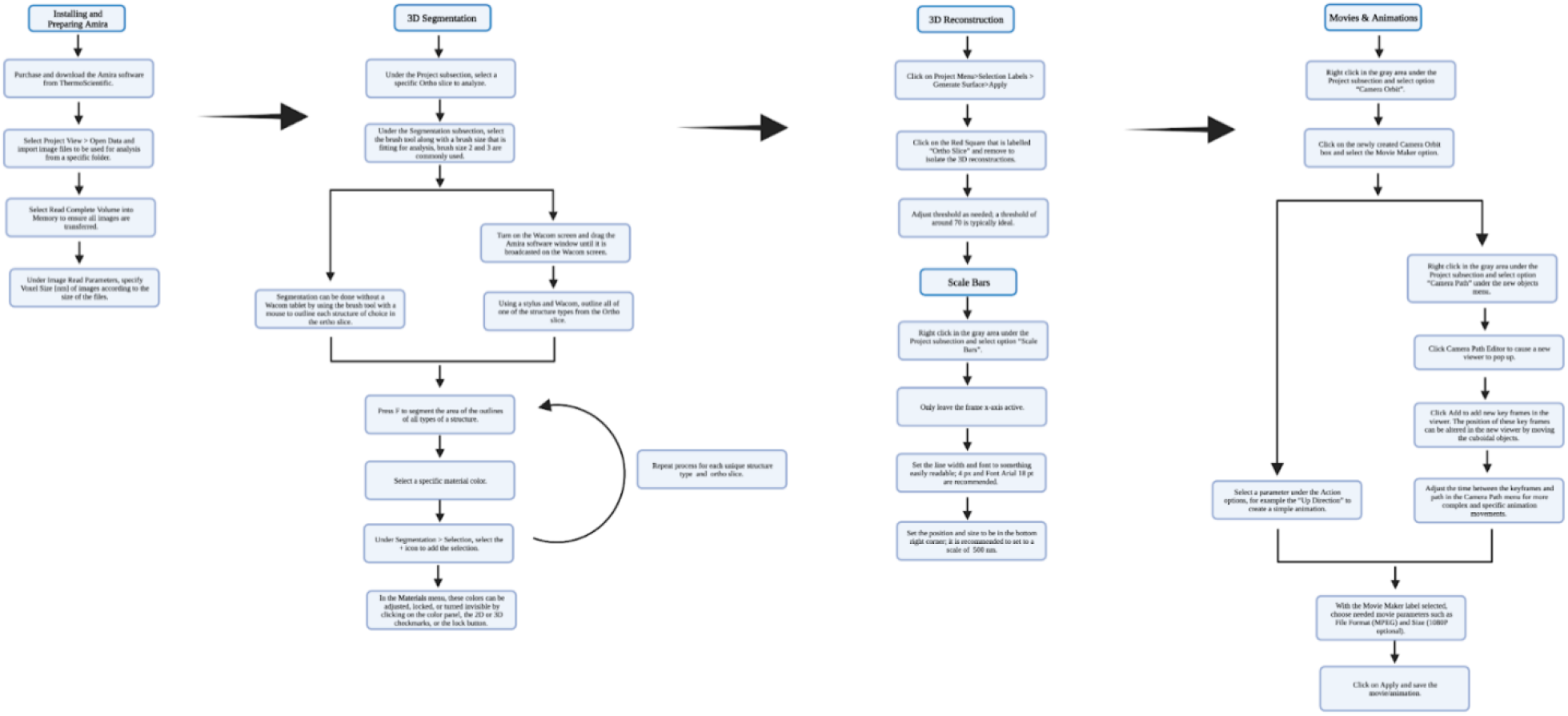
Printable flowchart of the optimized protocol for organelle quantification using Amira. **Description:** Standardized protocol for Amira segmentation, 3D reconstruction, and basic animation creation for organelles. In this process, a Wacom tablet allows for greater autonomy when mapping data elements. Segmenting each ortho slice individually allows for greater control over the process and ensures that high quality and precise 3D images will be generated. In this flowchart, Wacom tablets are used to outline mitochondria and tracheae; however, organelles of any shape or size can be segmented using the proper drawing tools. This flowchart highlights how Amira provides the freedom and flexibility to perform customized tasks and operations. This flowchart may be useful as lab reference material.

**Supplementary Figure 2:**
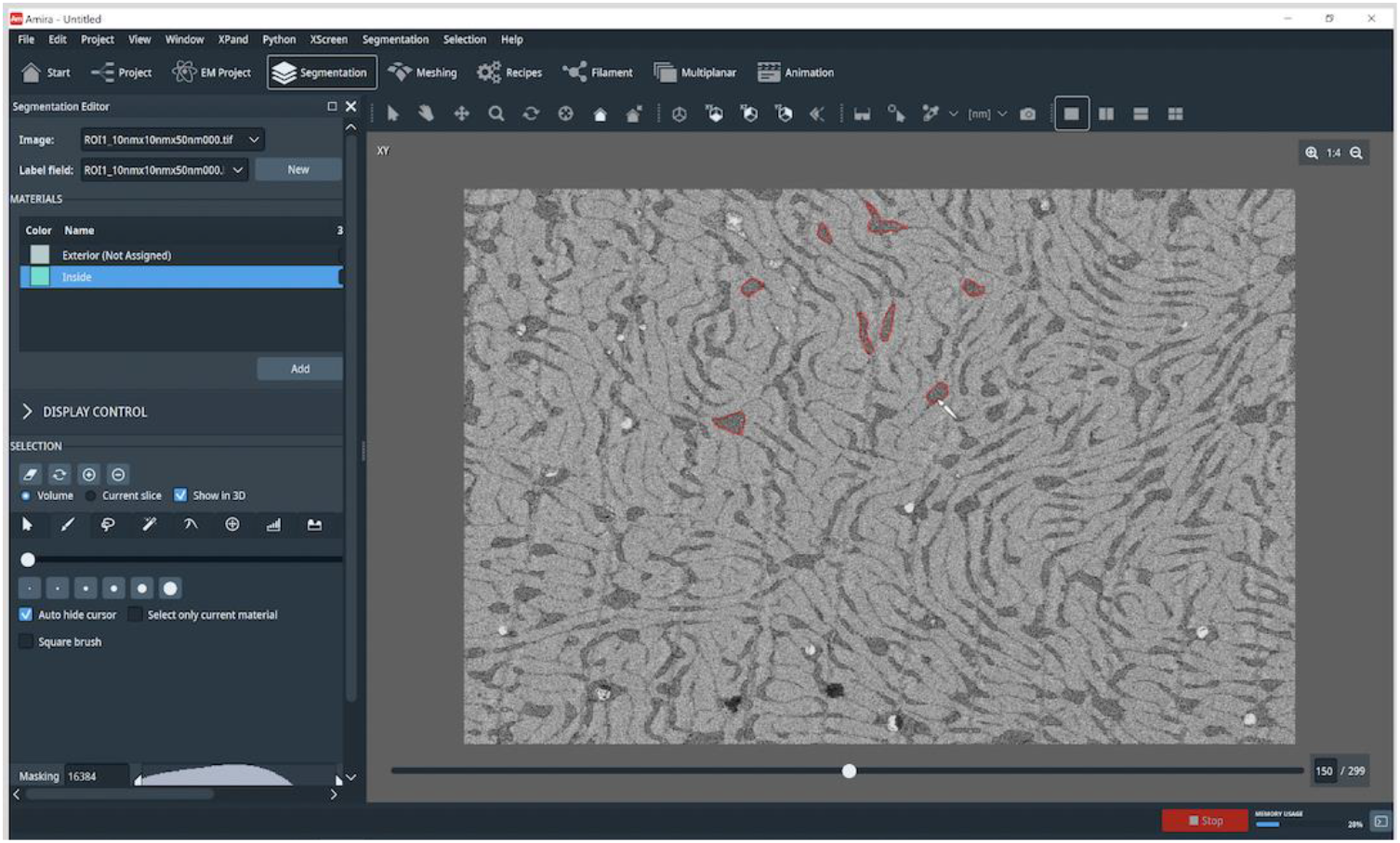
Sample of the Amira user interface. The segmentation tab on the Amira user interface. To the left is the main control interface that allows for control of the various sections. This methodology requires switching between the project, segmentation, and animation tabs. Under the segmentation section, the bottom left allows for custom control over the brush types and brush sizes. Under the display control subsection, color mapping and zooming may be altered, if relevant. Above the display control section is the materials subsection that allows for various colors, known as materials, to be selected to represent various organelles. In the materials subsection, colors can be modified, names can be changed, and 2D and 3D areas can be selected, adjusted, or locked. Additional materials for separate organelles can be added by selecting the Add button. The main area of the Amira interface contains the image to be analyzed. On this black and white SBF-SEM microscopy image, the highlighted areas show where tracheal segmentation has been performed. Above the image are various tools that allow for control over the function of the mouse, including a pointer, a 3D grabber, a color select tool, and a camera tool. The Project tab (not selected) is important for 3D rendering once segmentation has been completed.

## REFERENCES

1. Friederich, M.; Hansell, P.; Palm, F. Diabetes, Oxidative Stress, Nitric Oxide and Mitochondria Function. Current diabetes reviews 2009, 5, 120–144.

2. Bratic, A.; Larsson, N.-G. The Role of Mitochondria in Aging. The Journal of clinical investigation 2013, 123, 951–957.

3. Barja, G. The Mitochondrial Free Radical Theory of Aging. Progress in molecular biology and translational science 2014, 127, 1–27.

4. Olichon, A.; Baricault, L.; Gas, N.; Guillou, E.; Valette, A.; Belenguer, P.; Lenaers, G. Loss of OPA1 Perturbates the Mitochondrial Inner Membrane Structure and Integrity, Leading to Cytochrome c Release and Apoptosis. Journal of Biological Chemistry 2003, 278, 7743–7746.

5. Otera, H.; Miyata, N.; Kuge, O.; Mihara, K. Drp1-Dependent Mitochondrial Fission via MiD49/51 Is Essential for Apoptotic Cristae Remodeling. Journal of Cell Biology 2016, 212, 531–544.

6. Gunter, T.E.; Yule, D.I.; Gunter, K.K.; Eliseev, R.A.; Salter, J.D. Calcium and Mitochondria. FEBS letters 2004, 567, 96–102.

7. Dias, N.; Bailly, C. Drugs Targeting Mitochondrial Functions to Control Tumor Cell Growth. Biochemical pharmacology 2005, 70, 1–12.

8. Pinton, P.; Giorgi, C.; Siviero, R.; Zecchini, E.; Rizzuto, R. Calcium and Apoptosis: ER-Mitochondria Ca 2+ Transfer in the Control of Apoptosis. Oncogene 2008, 27, 6407–6418.

9. Nicholls, D.G. Mitochondria and Calcium Signaling. Cell calcium 2005, 38, 311–317.

10. Szewczyk, A.; Wojtczak, L. Mitochondria as a Pharmacological Target. Pharmacological reviews 2002, 54, 101–127.

11. Glancy, B. Visualizing Mitochondrial Form and Function within the Cell. Trends in molecular medicine 2020, 26, 58–70.

12. Transmission Electron Microscopy vs Scanning Electron Microscopy; ThermoScientific;

13. Arborgh, B.; Bell, P.; Brunk, U.; Collins, V. The Osmotic Effect of Glutaraldehyde during Fixation. A Transmission Electron Microscopy, Scanning Electron Microscopy and Cytochemical Study. Journal of ultrastructure research 1976, 56, 339–350.

14. Courson, J.A.; Landry, P.T.; Spehlmann, E.; Lafontant, P.; Patel, N.; Rumbaut, R.; Burns, A. Serial Block-Face Scanning Electron Microscopy (SBF-SEM) of Biological Tissue Samples. Journal of Visualized Experiments: Jove 2021.

15. Goggin, P.; Ho, E.M.; Gnaegi, H.; Searle, S.; Oreffo, R.O.; Schneider, P. Development of Protocols for the First Serial Block-Face Scanning Electron Microscopy (SBF SEM) Studies of Bone Tissue. Bone 2020, 131, 115107.

16. Baena, V.; Schalek, R.L.; Lichtman, J.W.; Terasaki, M. Serial-Section Electron Microscopy Using Automated Tape-Collecting Ultramicrotome (ATUM). Methods in cell biology 2019, 152, 41–67.

17. Denk, W.; Horstmann, H.; Harris, K.M. Serial Block-Face Scanning Electron Microscopy to Reconstruct Three-Dimensional Tissue Nanostructure. PLoS biology 2004, 2, e329.

18. Lippens, S.; Kremer, A.; Borghgraef, P.; Guérin, C.J. Serial Block Face-Scanning Electron Microscopy for Volume Electron Microscopy. Methods in cell biology 2019, 152, 69–85.

19. Hughes, L.; Hawes, C.; Monteith, S.; Vaughan, S. Serial Block Face Scanning Electron Microscopy—the Future of Cell Ultrastructure Imaging. Protoplasma 2014, 251, 395–401.

20. Mukherjee, K.; Clark, H.R.; Chavan, V.; Benson, E.K.; Kidd, G.J.; Srivastava, S. Analysis of Brain Mitochondria Using Serial Block-Face Scanning Electron Microscopy. Journal of visualized experiments: JoVE 2016.

21. Howard, V.; Reed, M. Unbiased Stereology: Three-Dimensional Measurement in Microscopy; Garland Science, 2004; ISBN 1-135-33168-5.

22. Lam, J.; Katti, P.; Biete, M.; Mungai, M.; AshShareef, S.; Neikirk, K.; Lopez, E.G.; Vue, Z.; Christensen, T.A.; Beasley, H.K. A Universal Approach to Analyzing Transmission Electron Microscopy with ImageJ. bioRxiv 2021.

23. Cogliati, S.; Enriquez, J.A.; Scorrano, L. Mitochondrial Cristae: Where Beauty Meets Functionality. Trends in biochemical sciences 2016, 41, 261–273.

24. Kühlbrandt, W. Structure and Function of Mitochondrial Membrane Protein Complexes. BMC biology 2015, 13, 1–11.

25. Wolf, S.G.; Mutsafi, Y.; Dadosh, T.; Ilani, T.; Lansky, Z.; Horowitz, B.; Rubin, S.; Elbaum, M.; Fass, D. 3D Visualization of Mitochondrial Solid-Phase Calcium Stores in Whole Cells. Elife 2017, 6, e29929.

26. Frey, T.G.; Mannella, C.A. The Internal Structure of Mitochondria. Trends in biochemical sciences 2000, 25, 319–324.

27. Jährling, N.; Becker, K.; Schönbauer, C.; Schnorrer, F.; Dodt, H.-U. Three-Dimensional Reconstruction and Segmentation of Intact Drosophila by Ultramicroscopy. Frontiers in systems neuroscience 2010, 4, 1.

28. Kopecky, B.; Duncan, J.; Elliott, K.; Fritzsch, B. Three-dimensional Reconstructions from Optical Sections of Thick Mouse Inner Ears Using Confocal Microscopy. Journal of microscopy 2012, 248, 292–298.

29. Jährling, N.; Becker, K.; Dodt, H.-U. 3D-Reconstruction of Blood Vessels by Ultramicroscopy. Organogenesis 2009, 5, 227–230.

30. Iwai, M.; Yokono, M.; Nakano, A. Visualizing Structural Dynamics of Thylakoid Membranes. Scientific reports 2014, 4, 1–6.

31. Cocks, E.; Taggart, M.; Rind, F.; White, K. A Guide to Analysis and Reconstruction of Serial Block Face Scanning Electron Microscopy Data. Journal of microscopy 2018, 270, 217–234.

32. Stalling, D.; Westerhoff, M.; Hege, H.-C. Amira: A Highly Interactive System for Visual Data Analysis. The visualization handbook 2005, 38, 749–67.

33. Suga, S.; Nakamura, K.; Humbel, B.M.; Kawai, H.; Hirabayashi, Y. An Interactive Deep Learning-Based Approach Reveals Mitochondrial Cristae Topologies. bioRxiv 2021.

34. Frezza, C.; Cipolat, S.; De Brito, O.M.; Micaroni, M.; Beznoussenko, G.V.; Rudka, T.; Bartoli, D.; Polishuck, R.S.; Danial, N.N.; De Strooper, B. OPA1 Controls Apoptotic Cristae Remodeling Independently from Mitochondrial Fusion. Cell 2006, 126, 177–189.

35. Olichon, A.; Guillou, E.; Delettre, C.; Landes, T.; Arnauné-Pelloquin, L.; Emorine, L.J.; Mils, V.; Daloyau, M.; Hamel, C.; Amati-Bonneau, P. Mitochondrial Dynamics and Disease, OPA1. Biochimica et Biophysica Acta (BBA)-Molecular Cell Research 2006, 1763, 500–509.

36. Sutherland, D.; Samakovlis, C.; Krasnow, M.A. Branchless Encodes a Drosophila FGF Homolog That Controls Tracheal Cell Migration and the Pattern of Branching. Cell 1996, 87, 1091–1101.

37. Sanz, A.; Stefanatos, R.K. The Mitochondrial Free Radical Theory of Aging: A Critical View. Current aging science 2008, 1, 10–21.

38. Rowland, A.A.; Voeltz, G.K. Endoplasmic Reticulum–Mitochondria Contacts: Function of the Junction. Nature reviews Molecular cell biology 2012, 13, 607–615.

39. Giacomello, M.; Pellegrini, L. The Coming of Age of the Mitochondria–ER Contact: A Matter of Thickness. Cell Death & Differentiation 2016, 23, 1417–1427.

40. Scorrano, L.; De Matteis, M.A.; Emr, S.; Giordano, F.; Hajnóczky, G.; Kornmann, B.; Lackner, L.L.; Levine, T.P.; Pellegrini, L.; Reinisch, K. Coming Together to Define Membrane Contact Sites. Nature communications 2019, 10, 1–11.

41. Kim, Y.; Lindberg, E.; Bleck, C.K.; Glancy, B. Endothelial Cell Nanotube Insertions into Cardiac and Skeletal Myocytes during Coordinated Tissue Development. Cardiovascular research 2020, 116, 260–261.

42. Merkwirth, C.; Langer, T. Mitofusin 2 Builds a Bridge between ER and Mitochondria. Cell 2008, 135, 1165–1167.

43. De Brito, O.M.; Scorrano, L. Mitofusin 2 Tethers Endoplasmic Reticulum to Mitochondria. Nature 2008, 456, 605–610.

44. Filadi, R.; Greotti, E.; Pizzo, P. Highlighting the Endoplasmic Reticulum-Mitochondria Connection: Focus on Mitofusin 2. Pharmacological research 2018, 128, 42–51.

45. Ngoh, G.A.; Papanicolaou, K.N.; Walsh, K. Loss of Mitofusin 2 Promotes Endoplasmic Reticulum Stress. Journal of Biological Chemistry 2012, 287, 20321–20332.

46. Yasukawa, K.; Oshiumi, H.; Takeda, M.; Ishihara, N.; Yanagi, Y.; Seya, T.; Kawabata, S.; Koshiba, T. Mitofusin 2 Inhibits Mitochondrial Antiviral Signaling. Science signaling 2009, 2, ra47–ra47.

47. Filadi, R.; Greotti, E.; Turacchio, G.; Luini, A.; Pozzan, T.; Pizzo, P. Mitofusin 2 Ablation Increases Endoplasmic Reticulum–Mitochondria Coupling. Proceedings of the National Academy of Sciences 2015, 112, E2174–E2181.

48. Naon, D.; Zaninello, M.; Giacomello, M.; Varanita, T.; Grespi, F.; Lakshminaranayan, S.; Serafini, A.; Semenzato, M.; Herkenne, S.; Hernández-Alvarez, M.I. Critical Reappraisal Confirms That Mitofusin 2 Is an Endoplasmic Reticulum–Mitochondria Tether. Proceedings of the National Academy of Sciences 2016, 113, 11249–11254.

49. Sebastián, D.; Hernández-Alvarez, M.I.; Segalés, J.; Sorianello, E.; Muñoz, J.P.; Sala, D.; Waget, A.; Liesa, M.; Paz, J.C.; Gopalacharyulu, P. Mitofusin 2 (Mfn2) Links Mitochondrial and Endoplasmic Reticulum Function with Insulin Signaling and Is Essential for Normal Glucose Homeostasis. Proceedings of the National Academy of Sciences 2012, 109, 5523–5528.

50. Sun, M.G.; Williams, J.; Munoz-Pinedo, C.; Perkins, G.A.; Brown, J.M.; Ellisman, M.H.; Green, D.R.; Frey, T.G. Correlated Three-Dimensional Light and Electron Microscopy Reveals Transformation of Mitochondria during Apoptosis. Nature cell biology 2007, 9, 1057–1065.

51. Peddie, C.J.; Collinson, L.M. Exploring the Third Dimension: Volume Electron Microscopy Comes of Age. Micron 2014, 61, 9–19.

52. Briggman, K.L.; Bock, D.D. Volume Electron Microscopy for Neuronal Circuit Reconstruction. Current opinion in neurobiology 2012, 22, 154–161.

53. Lidke, D.S.; Lidke, K.A. Advances in High-Resolution Imaging–Techniques for Three-Dimensional Imaging of Cellular Structures. Journal of cell science 2012, 125, 2571–2580.

54. Renken, C.W. The Structure of Mitochondria. 2004.

55. Willingham, T.B.; Kim, Y.; Lindberg, E.; Bleck, C.K.; Glancy, B. The Unified Myofibrillar Matrix for Force Generation in Muscle. Nature communications 2020, 11, 1–10.

56. Bleck, C.K.; Kim, Y.; Willingham, T.B.; Glancy, B. Subcellular Connectomic Analyses of Energy Networks in Striated Muscle. Nature communications 2018, 9, 1–11.

57. Vincent, A.E.; White, K.; Davey, T.; Philips, J.; Ogden, R.T.; Lawless, C.; Warren, C.; Hall, M.G.; Ng, Y.S.; Falkous, G. Quantitative 3D Mapping of the Human Skeletal Muscle Mitochondrial Network. Cell reports 2019, 26, 996–1009.

58. Faitg, J.; Lacefield, C.; Davey, T.; White, K.; Laws, R.; Kosmidis, S.; Reeve, A.K.; Kandel, E.R.; Vincent, A.E.; Picard, M. 3D Neuronal Mitochondrial Morphology in Axons, Dendrites, and Somata of the Aging Mouse Hippocampus. bioRxiv 2021.

59. Soufan, A.T.; Ruijter, J.M.; van den Hoff, M.J.; de Boer, P.A.; Hagoort, J.; Moorman, A.F. Three-Dimensional Reconstruction of Gene Expression Patterns during Cardiac Development. Physiological genomics 2003, 13, 187–195.

60. de Boer, B.A.; Soufan, A.T.; Hagoort, J.; Mohun, T.J.; van den Hoff, M.J.; Hasman, A.; Voorbraak, F.P.; Moorman, A.F.; Ruijter, J.M. The Interactive Presentation of 3D Information Obtained from Reconstructed Datasets and 3D Placement of Single Histological Sections with the 3D Portable Document Format. Development 2011, 138, 159–167.

61. Lanika Solutions Amira Software For Life & Biomedical Sciences; ThermoScientific;

62. Schneider, C.A.; Rasband, W.S.; Eliceiri, K.W. NIH Image to ImageJ: 25 Years of Image Analysis. Nature methods 2012, 9, 671–675.

63. Berg, S.; Kutra, D.; Kroeger, T.; Straehle, C.N.; Kausler, B.X.; Haubold, C.; Schiegg, M.; Ales, J.; Beier, T.; Rudy, M. Ilastik: Interactive Machine Learning for (Bio) Image Analysis. Nature Methods 2019, 16, 1226–1232.

64. Mustafi, D.; Kikano, S.; Palczewski, K. Serial Block Face-Scanning Electron Microscopy: A Method to Study Retinal Degenerative Phenotypes. Current protocols in mouse biology 2014, 4, 197–204.

